# Genome-wide assessment of REST binding profiles reveals distinctions between human and mouse hippocampus

**DOI:** 10.1101/2020.07.07.192229

**Authors:** James C. McGann, Michael Spinner, Saurabh K. Garg, Karin Mullendorf, Randall L. Woltjer, Gail Mandel

**Affiliations:** Vollum Institute, Oregon Health and Science University, Portland, OR, USA; Cancer Early Detection Advanced Research Center, Oregon Health and Science Institute, Portland, OR, USA; Department of Cutaneous Oncology, H. Lee Moffitt Cancer Center and Research Institute, Tampa, FL, USA; Universidad San Sebastian, Puerto Montt, Chile; Department of Pathology, Division of Neuropathology, Oregon Health and Science University, Portland, OR, USA

## Abstract

**Background:** The transcriptional repressor, RE1 Silencing Transcription Factor (REST), recognized historically as a master regulator of neuronal gene expression during mouse development, has recently been ascribed roles in human aging and neurodegenerative disorders. However, REST’s role in healthy adult human brain, and how faithfully mouse models reproduce REST function in human brain, is not known.

**Results:** Here, we present the first genome-wide binding profile for REST in both mouse and human postnatal hippocampus. We find the majority of REST-bound sites in human hippocampus are unique compared to both mouse hippocampus and to all other reported human ENCODE cell types. Genes associated with unique REST-bound sites include previously unidentified categories related to innate immunity and inflammation signaling, suggesting species specific roles for REST in protecting human brain health.

**Conclusions:** Our results suggest newly evolved functions for REST in maintaining human brain health.

## Introduction

The role of transcription factors during the development and early maturation of the nervous system has been the subject of intense research, as new sequencing technologies have been brought to bear on neurodevelopmental disorders such as autism and schizophrenia. A factor of interest in this regard is the RE1 Silencing Transcription Factor (REST [1]; also called NRSF [2]). In vertebrates, REST is expressed in non-neuronal cells outside and within the nervous system, in non-neuronal embryonic stem cells and in neural progenitors. In the non-neuronal cells, REST binds to thousands of target genes, both coding and non-coding, containing its consensus binding site, the 21-nucleotide Repressor Element 1 (RE1 [3]; NRSE [4,5,6]). The relief of REST-mediated repression including dismissal of its co-repressors during differentiation [7], endows mature phenotypic features on neurons, amplifying the difference between neuronal and non-neuronal cells [8]. Neuronal features regulated by REST include voltage-sensitive and ligand-gated ion channels, growth factors and their cognate receptors, synaptic proteins, and neuronal transcription factors (reviewed in Ballas and Mandel, 2005 [9]).

Historically, most work on REST has focused exclusively on its important role in forming the rodent embryonic nervous system. More recent studies, however, indicate that REST also plays a role later in life. For example, several studies in rodents indicate that following terminal neuronal differentiation, REST levels increase from minimal to significantly higher levels in certain populations of post-mitotic neurons in late embryogenesis, persisting in the postnatal brain [10,11,12,13,14,15]. REST levels are also increased in conditions of ischemia or seizure in adult mouse brain, and preventing the increase alleviates neuronal cell death associated with both conditions [12,16,17,18,19,20]. Regarding human REST, increased levels are reported to provide protective roles during aging through binding to genes associated with cell death pathways and nervous system excitability [21,22].

Several additional molecular and bioinformatics findings motivate this study. First, while REST initially appeared with the evolution of a vertebrate nervous system [23], REST has also evolved with strong positive selection in humans specifically [24,25]. Second, comparisons of REST binding profiles in human and mouse embryonic stem cells, and across other human cell types, have indicated an expansion of the human cistrome, suggesting unique roles for REST-regulated genes in human brain development and neurodevelopmental disease [26,27,28,29]. Third, the cognate RE1 sequence is also highly conserved across species [23], although many RE1s have evolved to be unique to the human genome [27,29]. These genome-wide findings notwithstanding, whether the human brain reflects a more expansive cistrome than mouse brain, commensurate with the higher functional and anatomical complexity in human brain, has not been addressed. Here, we address this question using chromatin immunoprecipitation with massively parallel DNA sequencing (ChIP-seq) on mouse and human hippocampus. We focus on the hippocampus, specifically, because of previous work indicating a role for REST in aging and cognition [21], and our work herein, indicating dichotomous REST developmental expression patterns in human and mouse hippocampus.

## Results

### REST-bound sites in the mouse postnatal hippocampus

REST target genes have not been identified genome-wide by ChIP-seq in either the murine or human brain. We focused on elucidating the mouse REST-bound targets first because most experimental work on REST relied on murine models and murine cell lines, and because of the advantage of REST KO cells for validating REST function. To this end, we compared the REST-bound peaks in hippocampus with peaks found previously in Embryonic Stem Cells (ESCs) and the C2C12 muscle cell line (Fig. 1a, [30,31]). We performed the ChIP-seq analysis on 5w old mice because previous studies, and the study herein, indicated that REST was expressed to moderately high levels in brain regions of postnatal mice at 5w age (Additional file 1: Figure S1.1a-d). For the REST immunoprecipitation, we used an affinity-purified antibody generated by our laboratory raised against the C-terminus of full-length human REST. Western blot immuno-labeling and qChIP in *Rest* knockout mice confirmed specificity of the antibody to full length REST protein [15] (Additional file 1: Figure S1.1d-e). Of note, this antibody will not recognize truncated splice variants [32], whose functional importance and expression levels are still characterized poorly and are not considered in this study.

**Figure 1.**
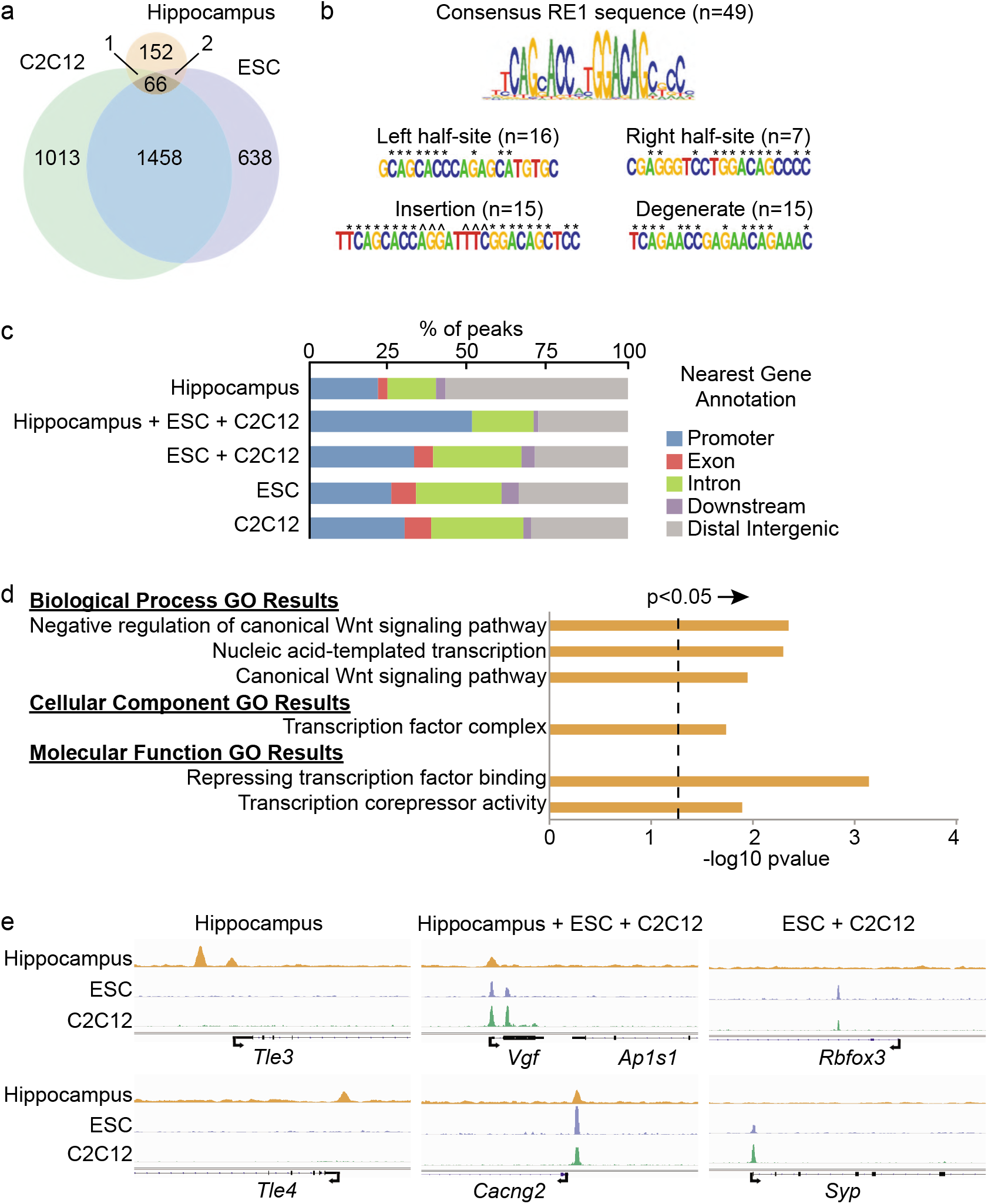
REST ChIP-seq analysis reveals a new set of associated genes in mouse hippocampus. a) Venn diagram showing number of REST-bound peaks and overlap among hippocampus and each of two non-neuronal cell types. b) Consensus RE1 sequences and related RE1 sequences. Values in parenthesis denote the number of hippocampal REST ChIP-seq peaks. * denotes nucleotides that match the RE1 consensus, ^ denotes inserted nucleotides. c) Distribution of REST peaks at RE1 consensus and related sequences relative to the nearest annotated gene. d) Gene ontology categories for genes associated with REST peaks unique to the hippocampus. Negative log10 of p-value was calculated by Fisher’s exact test and corrected by False Discovery Rate (Methods). E) REST peaks at representative genomic loci. Tle loci are unique to hippocampus and other loci indicated are prototypical neuronal RE1 sequences. Note: all REST peaks shown have underlying RE1 or related sequences with the exception of Tle3. Arrows denote transcriptional start sites. Broader horizontal lines in genes represent exons.

We identified only 221 REST-bound peaks in the mouse hippocampus, with only 30% of the peaks also shared with C2C12 and ESCs (Fig. 1a). Heat maps further reflected the reduced overlap, as REST densities in the peaks were much reduced or absent in the other cell types (Fig. S1.2). The paucity of REST peaks in hippocampus likely reflects the lower REST expression levels compared to ESCs and non-neuronal cells. Additionally, REST levels generally drop as differentiation proceeds [7,33]. We sought to determine whether the unique REST-bound hippocampal peaks were associated with genes lacking consensus and related RE1 sequences typical of nonneuronal cells and neural progenitors, since binding to non-RE1 sequences has been reported previously [27,28]. For this analysis, we used consensus RE1 motifs, RE1 half motifs and other RE1-related motifs [5,6,23] (Fig. 1b). As expected, all hippocampal peaks shared with non-neuronal cells displayed consensus or RE1-related sequences. However, of the REST peaks unique to the hippocampus, none overlaid a consensus RE1 sequence and only 22% were associated with an RE1-related sequence (Table 1; Additional file 2: Table S1). In contrast, consensus and RE1-related peaks represented 88% and 82% respectively of the sites unique to each non-neuronal cell type (Table 1). The distribution of REST peaks shared among all three tissue/cell samples, which contained RE1 sequences, was biased more towards the promoter regions of annotated genes (Fig. 1c), consistent with previous studies of RE1 locations in mouse ESCs [27,30]. The distribution of REST peaks unique to hippocampus was biased away from promoters and more towards intergenic regions. More than 50% of the hippocampal-unique peaks belonged to this category compared to only ~25% intergenic peaks shared among all cell types (Fig. 1c), commensurate with previous findings that nonconserved sites are located more distally from promoters [29].

**Table 1.** REST peak number and underlying RE1 motif status for mouse hippocampus, mESCs and mC2C12 myoblasts

| Tissue/Cell type | With consensus RE1 | With related RE1 | Lack any RE1 |
| --- | --- | --- | --- |
| Hippocampal unique (152) | 0 | 33 | 119 |
| Hippocampal, ESC, C2C12, shared (66) | 48 | 18 | 0 |
| ESC, C2C12, shared (1458) | 1105 | 351 | 2 |
| ESC unique (638) | 292 | 271 | 75 |
| C2C12 unique (1013) | 331 | 497 | 185 |

Summary of ChIP-seq results comparing REST binding characteristics among hippocampus and C2C12 muscle cells and ESCs. m refers to mouse. Numbers in parentheses refer to total peak numbers

Because of the presence of binding sites lacking RE1 sequences, we considered the possibility that the associated target genes in hippocampus might represent a distinct gene set. To test this idea, we first defined REST target genes as annotated genes with REST peaks in the gene body extending from 5kb upstream of the Refseq-annotated transcriptional start site (TSS) to 3kb downstream of the 3’untranslated regions. For the REST peaks shared among hippocampus and the non-neuronal cells, all of which contained RE1 or RE1-related sequences, the Gene Ontology (GO) enrichments were very similar to those from either ESCs or C2C12 cells (Additional file 2: Table S2) and neural progenitors [7]. These GO categories represented prototypical REST-regulated genes important in neurodevelopment, such as those related to glutamatergic synaptic transmission (33-fold enrichment, GO:0035249), neurotransmitter binding (14-fold enrichment, GO:0042165), and postsynaptic membrane localization (GO:0045211). In contrast to these shared “neuronal” REST target genes, supported by many experimental studies, GO terms for the target genes associated with unique hippocampal REST peaks showed enrichments for less stereotypical functions (Fig. 1d,e). For example, transcription factors were a highly enriched category. Interestingly, several paralogs of the transducing-like enhancer of split family of transcriptional co-repressors (*Tle1, Tle3, Tle4*, and *Tle6*) accounted for this category (Fig. 1e). This finding is consistent with a previous study indicating that a single GO term can reach significance due to the presence of highly related genes in a gene family [23]. Both *Tle1* and *Tle4* had RE1-related sequences, while *Tle3* and *Tle6* had no discernable RE1 site. The sequences underlying these, as well as the other unique hippocampal peaks lacking RE1 or related sequence, did not show any enriched binding motifs for other transcription factors (DIVERSITY algorithm [34]). The *Tle2* gene has an RE1 sequence, but was not occupied by REST in our ChIP-seq analysis.

Conversely, we also identified REST peaks coinciding with RE1 sequences in ESCs and C2C12 cells that were absent from hippocampus (Fig. 1a,e; Additional file 1: Table S1). Representative genes associated with these peaks included the *Hes3, Syp*, and *Nrxn1* genes that are enriched in similar neuronal GO categories (Additional file 2: Table S2). As REST and the underlying RE1 sequences are present in both neurons and glia in hippocampus, the factor(s) that prevent REST binding to these cognate sites, as well to the RE1 site associated with Tle2 gene described above, opens up an important problem for the future.

### REST-bound sites in human hippocampus

To first ensure that postnatal human hippocampus had sufficient REST protein levels for ChIP-seq analysis, we performed Western blot analysis (Additional file 1: Figure S1.3a). In human adult post mortem hippocampus, in contrast to canonical low levels of REST during embryogenesis, REST levels were moderately high from ~age 40 up until our latest point of analysis at 85y of age. The prominent REST expression in adult human brain was consistent with previous studies [21, 25]. Therefore, here, we focus our analysis on REST function in the age interval of 60-70 years when REST levels seemed sufficient for ChIP-seq analysis. Surprisingly, mice showed the opposite trend, with REST levels moderately high at the time of our ChIP-seq analysis in the juvenile mice (5w of age), but, unexpectedly, dropping dramatically to barely detectable levels by 4-6 months after birth (Additional file 1: Figure S1.3b). The reciprocal REST expression patterns between mouse and human hippocampus suggested different roles for REST, underscoring our interest in determining the REST target genes in human brain.

Unexpectedly, upon our initial investigation of the REST binding profiles, we found that a preponderance of the REST peaks coincided with nearest genes that were highly expressed and encoded ubiquitous proteins. Further scrutiny showed that these REST peaks coincided with High Occupancy Target (HOT) regions, also called phantom or hyper-ChIPable peaks identified in previous ChIP-seq analyses for transcription factors other than REST [35,36,37,38,39]. HOT regions in human, mouse, worm, and fly were all in the 99^th^ percentile of genomic regions for the combined density of transcription factor ChIP-seq enrichment from the ENCODE database [35], indicating that they were not specific for the factors. We used similar criteria for assessing whether our unusual REST peaks could be assigned to a HOT region category. We found that 1) REST peaks at putative HOT regions were present at TSSs as defined by ChIP-seq analyses for RNA polymerase II and chromatin markers of active gene expression (Additional file 3: Figure S1a), 2) REST peaks were present at HOT regions even in a control ChIP-seq using IgG for the pull down (Additional file 3: Figure S1a), 3) the presence of REST peaks at HOT regions was independent of the ENCODE cell type or REST antibody used for the ChIP analyses (Additional file 3: Figure S1a, b), and 4) REST peaks occurred at HOT regions even in a ChIP-seq analysis using an antibody to GFP against eGFP-tagged REST protein (Additional file 3: Figure S1b).

Authentic genomic targets of a specific transcription factor were identified previously by filtering out hyper-ChIPable peaks using a cell or animal model deleted for the transcription factor [39]. Because our analysis used postmortem brain tissue, we used a bioinformatics filter for our data (see methods). In brief, we used the aggregate plots for ChIP-seq read depths for H3K4me3 and H3K9ac modifications as a proxy for the HOT regions in the ENCODE dataset and compared these read depths to the level of REST enrichment at hippocampal sites (bigwigAverageOverBed, Additional file 3: Figure S1c). We then set a threshold that removed the majority of known HOT regions while maintaining REST peaks with RE1 and RE1-related sites (Additional file 3: Figure S1d, e). Our subsequent analyses of hippocampal REST ChIP-seq data included only the REST peaks that remained after applying the filter (Additional file 2: Table S3).

We identified a total of 2125 hippocampal REST peaks after filtering, 78% of which had a corresponding sequence in the mouse genome. Of the total peaks, 89% were unique relative to previously annotated REST peaks defined as REST-bound peaks in >40 of 48 ENCODE replicates from human non-neuronal cell lines (“Core” peaks; Fig. 2a; see Methods), consistent with a previous study indicating that REST occupancy is cell-specific [40]. Further, heat maps of REST ChIP densities showed that the full set of hippocampal peaks had distinct signatures among the publicly available REST ChIP-seq datasets (Additional file 1: Figure S2.1). The unique hippocampal REST peaks were associated with a much higher percentage of peaks lacking underlying consensus and related RE1 sequences compared to the Core REST peaks (38% versus 3%, respectively; Table 2). The genomic loci shared between hippocampus and Core were both biased more towards promoter regions (Fig. 2b). Predictably, the genes associated with the shared peaks, exemplified in HeLa cells and hESCs, were enriched for the typical neuronal GO categories including growth factors, ion channels, neurotransmitter receptors and synaptic adhesion molecules (Fig. 2c; Additional file 2; Table S4). We noted that REST enrichment in the hippocampal unique peaks was overall lower than REST enrichment in the peaks shared with the Core dataset (Fig. 2c, Additional file 1: Figure S3.1a). However, lower levels of REST enrichment were also observed for cellspecific peaks in ESCs and A549 cells when each was compared to the other cell types in the Core dataset (Additional file 1: Figure S3.1b), and is consistent with another study indicating that more ancient, i.e. conserved RE1 sites, have higher affinities than lineage-specific sites [29].

**Table 2.**
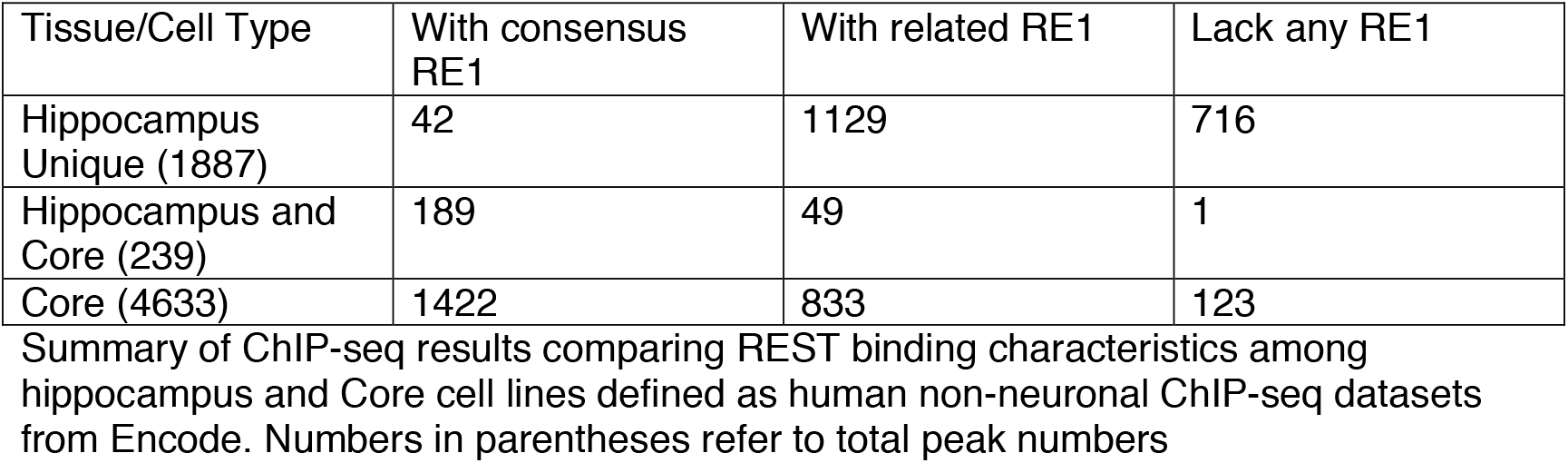
Human REST peak number and underlying RE1 motif status for hippocampus and Core cell lines

| Tissue/Cell Type | With consensus RE1 | With related RE1 | Lack any RE1 |
| --- | --- | --- | --- |
| Hippocampus Unique (1887) | 42 | 1129 | 716 |
| Hippocampus and Core (239) | 189 | 49 | 1 |
| Core (4633) | 1422 | 833 | 123 |

Summary of ChIP-seq results comparing REST binding characteristics among hippocampus and Core cell lines defined as human non-neuronal ChIP-seq datasets from Encode. Numbers in parentheses refer to total peak numbers

### The unique REST peaks in human hippocampus are largely absent from the same loci in mouse

Our ChIP-seq analysis indicated that the REST binding profile, in addition to being distinct from profiles in other human cell types, was also strikingly dissimilar from mouse hippocampal binding profiles (Additional file 2: Table S3). First, the total number of REST peaks in human hippocampus far exceeded the number of REST peaks in murine hippocampus (2125 versus 221 peaks, Figures 2a and 1a, respectively). This result is likely related to a lower hippocampal REST protein level in mouse compared to human (Additional file 2: Figure 3a,b). Second, only nine of the 1886 peaks unique to the human hippocampus (0.5%) were found at the corresponding location in mouse hippocampus or any murine non-neuronal cell type. Third, unlike unique peaks of mouse hippocampal ChIP, human unique peaks did not show bias towards intergenic regions (Fig. 1c and Fig. 2b). Conversely, when analyzing the complete dataset of REST target genes unique to the mouse hippocampus (Additional file 2: Table S1), none of these genes were associated with REST peaks in human hippocampus, and only one, *Gtf3c3* (General transcription factor 3c polypeptide 3), a gene involved in neurodevelopmental disorders [41], was also bound by REST in any of the human cell lines analyzed [31].

**Figure 2.**
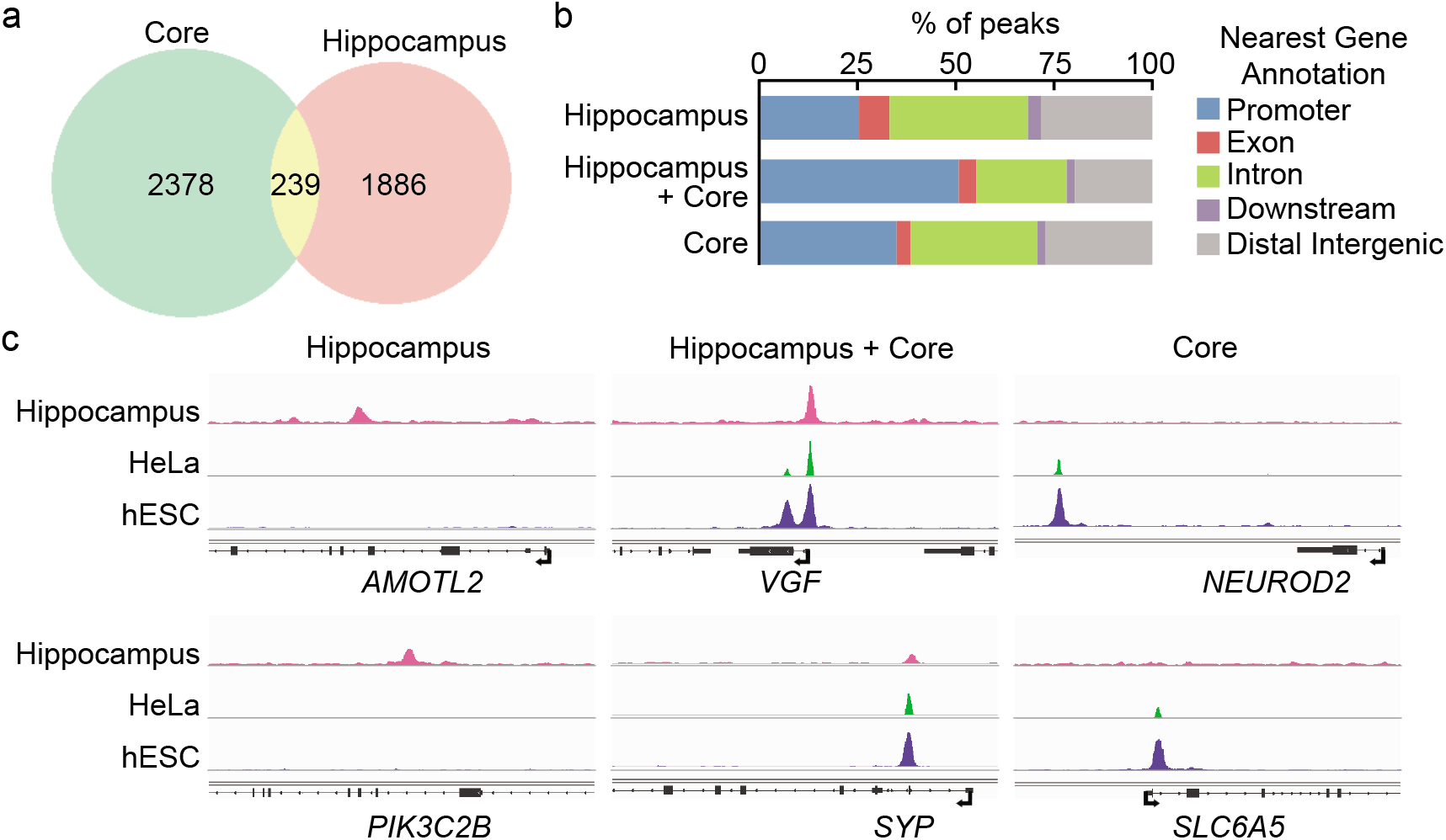
ChIP-seq analysis of human hippocampus. a) Venn diagram showing number of REST ChIP-seq peaks unique to the hippocampus and overlap with Core human REST peaks. Core peaks are defined as REST-bound peaks in >40 of 48 ENCODE replicates from human non-neuronal cell lines (Additional file 2: Table S5).

**Figure 3.**
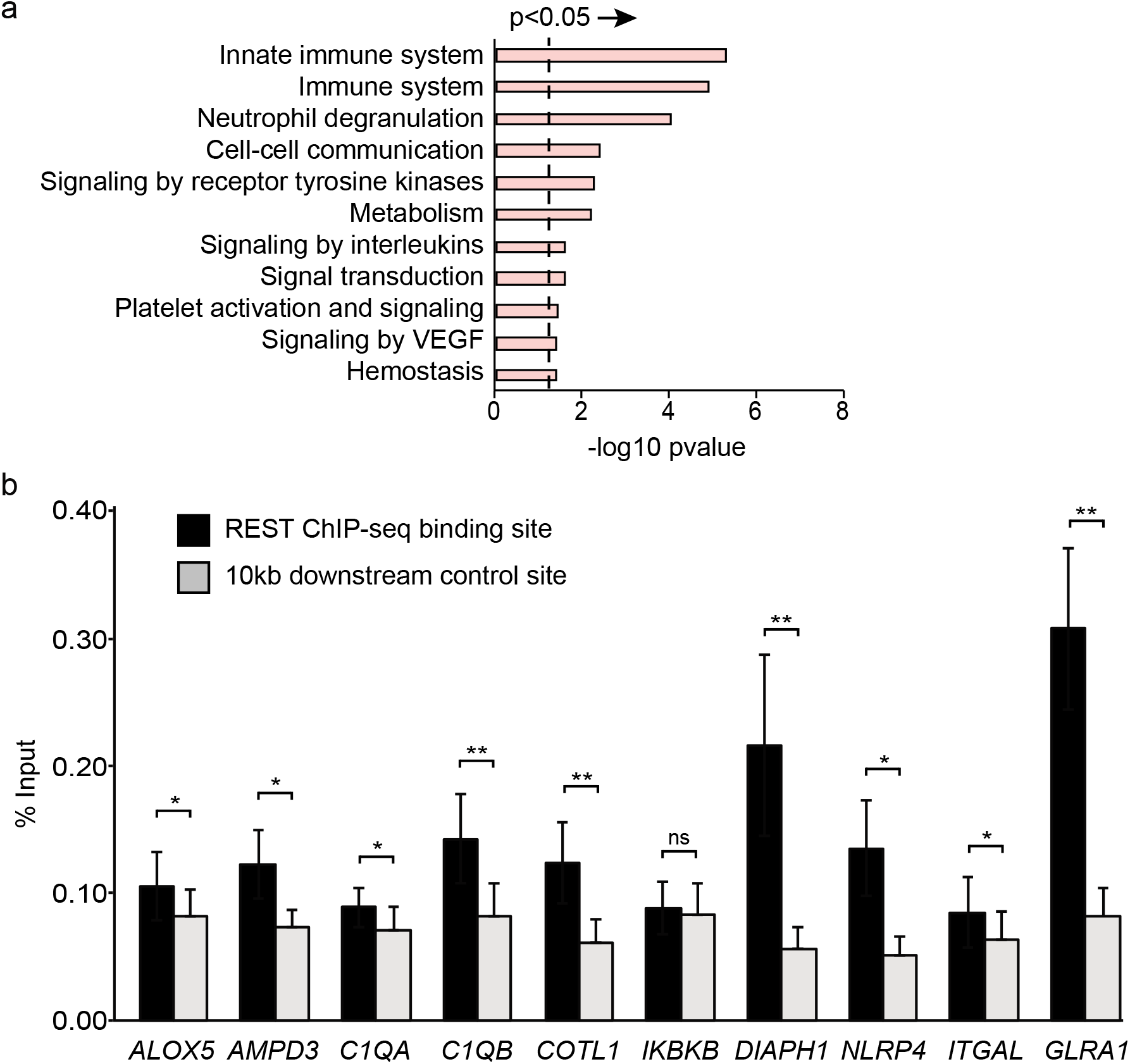
Human hippocampal-specific REST peaks are enriched near genes encoding proteins of non-canonical neuronal functions. a) Significant Reactome enrichment categories for genes associated with REST peaks unique to the hippocampus. p-value was calculated by Fisher’s exact test and corrected by False Discovery Rate (FDR, set to 0.05). b) Quantitative ChIP validation for a subset of the immune-related genes identified in the human hippocampal dataset using independent samples (n=8, 4 female and 4 male, ages 57-76yrs) * p<0.05, ** p<0.01, and **** p<0.0001 according to paired t-test test. The genes Alox5, Ampd3, C1qa, Cotl1, Ikbkb, Diaph1, and Itgal have degenerate RE1 sequences; the other two genes lack any discernable RE1.

### The Reactome categories of genes associated with REST hippocampal peaks include new categories related to the innate immunity system and signals of inflammation

We analyzed the unique hippocampal REST-associated genes by GO and Reactome category enrichments [42] (Additional file 2: Table S4). Surprisingly, in contrast to the REST-associated genes shared with the Core dataset, which were largely prototypical neuronal REST target genes, the top three Reactome enrichment categories included genes encoding proteins related to innate immunity (Fig. 3a). Additionally, enrichments in broad categories of signaling, including signaling in inflammatory processes (interleukins), as well as metabolism and hemostasis reached significance (Fig. 3a and Additional file 2: Table S4, p<0.05, Fisher’s exact test corrected for FDR of 0.05). We tested some of these genes by quantitative ChIP (qChIP) on hippocampal tissue acquired from similarly-aged individuals different from the tissue used previously for the ChIP-seq. A canonical REST target gene (*GLRA1*) with a consensus RE1 sequence served as a positive qChIP control. Primers for downstream sequences that lacked REST peaks served as negative controls for the REST antibody pull down. All but one of the ChIP-seq identified genes, *Inhibitor of Nuclear Factor Kappa B Kinase subunit Beta* (*IKBKB*), was validated by the qChIP results (Fig. 3b).

Finally, we sought to identify binding motifs for any other transcription factors that might recruit REST indirectly to the REST-bound sites, since 38% (716) of the unique human hippocampal peaks lacked RE1 or related sequences (Table 2). For this purpose, we applied the DIVERSITY program [34] to all sequences underlying the hippocampal REST peaks. In addition to the conserved RE1 motifs identified previously (Fig. 1b), DIVERSITY identified three enriched motifs related to known transcription factor binding motifs (Fig. 4). Notably, the predominant motif, detected in 663 peaks, significantly matched the motifs for the factors Sfpi1/PU.1, Prdm1, Irf1, and Etv6, as determined by a motif comparison tool, Tomtom [43] (FDR-adjusted p<0.05). These transcription factors are all involved in the regulation of immune cell development and function [44,45,46,47], consistent with the Reactome enrichments. Further, the two other identified motifs matched the binding sites for the transcription factors Sp2, Runx1, and Gfi1b, which regulate immune system differentiation [48,49,50] (Fig. 4). It seems likely that these factors are in complexes with REST, recruiting REST to the peak loci indirectly to mediate repression. This interpretation is the simplest and is supported by previous studies indicating the contribution of the REST co-repressor CoREST to some of these complexes [51]. For example, PU.1, Runx1 and Gfi1b, identified in our motif analysis, are in complexes with CoREST [52,53]. Further experiments confirming that REST is in complexes with the immune transcription factors in hippocampus are required for confirmation of this hypothesis.

**Figure 4.**
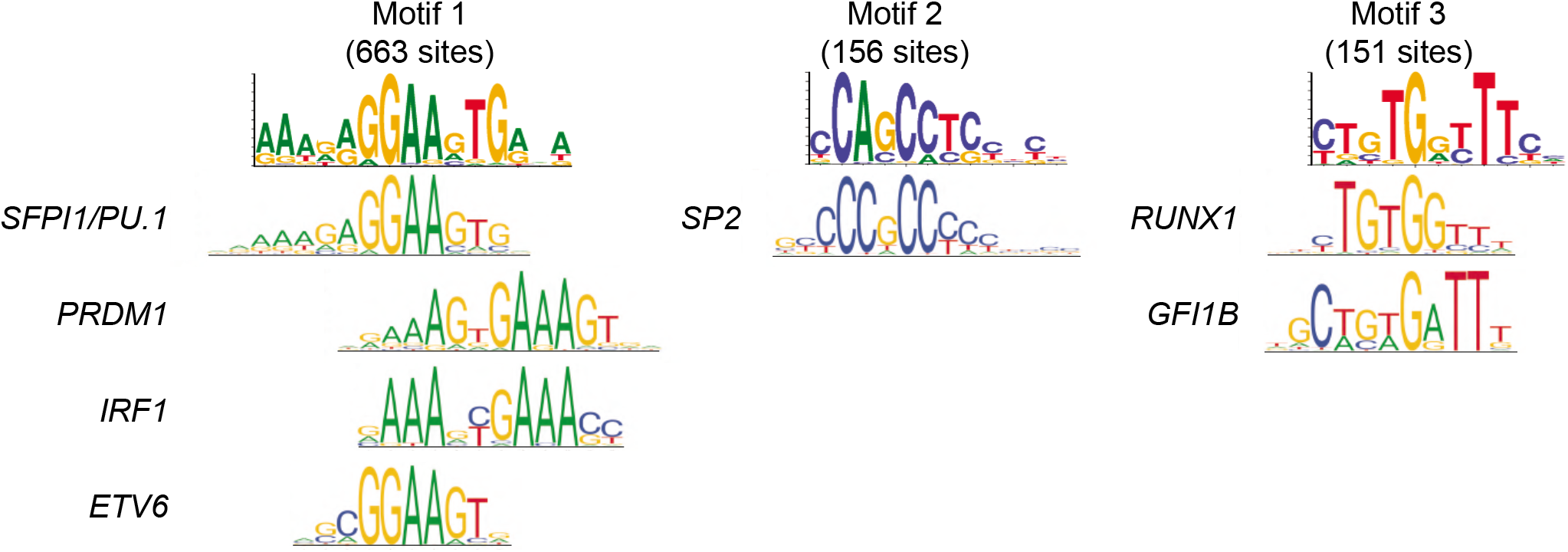
Non-RE1 motifs present in the REST peak dataset unique to the human hippocampus. Motifs were identified by DIVERSITY de novo motif analysis. The consensus motifs found in REST peaks is at the top and transcription factors that bind to related sequences are shown below. The number of sites refers to the number of REST peaks that are associated with a motif. Motifs were based on Tomtom motif analysis [43]. FDR-adjusted p<0.05.

## Discussion

Here, we have compared, for the first time, the genome-wide REST binding profiles in healthy post-natal mouse and human hippocampal chromatin at ages where REST levels were amenable to ChIP-seq analysis. We found that mouse and human REST hippocampus were highly dissimilar in terms of REST abundance at older ages, in the number of REST peaks, and in the nature of associated genes, raising questions about how well mouse postnatal development and aging would model the same processes in human brain. In particular, we have identified new gene classes associated with REST binding that are unique to the human hippocampus, representing largely, but not exclusively, processes related to innate immunity and inflammatory signaling.

We found that REST levels in mice declined to minimal levels in hippocampus at 4-6m, a time period associated with full brain maturity. Our REST-binding analysis, performed in 5-week old juvenile mice, prior to brain maturity, is most consistent with the historic model of REST. In this model, REST is still bound to its canonical neuronal genes in non-neuronal cells, progenitors, and immature neurons, awaiting its release from neuronal chromatin. At 4-6m, robust neurodevelopment is declining and a neurodevelopmental role for REST will be declining in parallel. While at least a fraction of the 152 unique hippocampal REST ChIP signals could be derived from neurons based on co-staining of REST with the neuronal marker MAP2 (Additional file 1: Figure S1.1b,c), previous studies indicate that most of the neurons may still be in an immature state [11,15]. This interpretation is consistent with REST-bound *Tle genes*. Not all paralogs of this family have been studied in mouse brain, but at least two of them function in regulating early neurodevelopment [54,55].

For our human REST ChIP-seq analysis, we filtered out previously reported HOT regions from our final analysis. To our knowledge, we are the only lab to have directly compared HOT regions in two different species using the same ChIP-seq antibody and experimental conditions. The basis for the human HOT-like REST peaks is not known, but one possibility we considered was that they represented looping between the promoter region and REST occupancy on distal sequences. This possibility seems unlikely because two independent ENCODE replicates of hESC and A451 ChIP-seq datasets differed greatly in the number of HOT REST peaks. Further, like the human genome, the mouse genome also has REST-bound sites located quite distal from TSSs [30], yet we did not find HOT regions in the ChIP-seq analysis of mouse hippocampus. Regarding this species difference, it is possible that the REST antibody is recognizing proteins non-specifically at the promoter regions of human highly expressed genes that are more accessible than promoters in mouse. While one report did note a trend for HOT region correlation with genomic accessibility [35], genomic accessibility did not appear to be sufficient, and we did not directly measure DNAse hypersensitivity at the HOT promoter regions bound by REST. In the future, confirmation of REST binding sites using alternative methods to measure REST binding *in vivo*, such as Cut and Run and Mnase-SSP [56,57], may shed more light on the curious human REST binding profile at some promoters.

Similar to findings from previous studies, a large percentage (38%) of REST peaks were associated with sequences not related to any variation of the RE1 sequence (Table 2; [27, 29]). Additionally, of the human hippocampal REST peaks that had no orthologous mouse sequence (462 peaks), a large percentage of them (54%) were associated with both consensus and non-consensus RE1 sequences. This result suggests that newly evolved RE1 sequences account for a large proportion of REST binding site expansion in human, and adds to previous results in hESCs where consensus RE1 sequences with non-conserved REST peaks were primate or human-specific [27,29].

REST has been studied experimentally from the perspective of human aging and age-related disease, particularly in the context of Alzheimer’s Disease (AD) [21]. For example, these investigators reported that REST levels continue to increase with human age, in neurons in both cortex and hippocampus, while levels decreased in AD tissue. Here, we confirm moderately high REST levels in intact human hippocampus at ~40-85 years of age, but we were unable to conclude there was a strong correlation with age, perhaps requiring analysis of a larger dataset (Additional file 1: Figure S1.3a). Because this is not an aging study, this experiment was not followed up here. Interestingly, in their study, Lu et al. further identified a set of REST repressed genes in neurons encoding proteins that promoted apoptosis and AD pathology, concentrating primarily on human prefrontal cortex, but also showing a correlation of REST levels and AD pathology in hippocampus as well. In our ChIP-seq analysis of postmortem hippocampus, although we failed to identify the same pro-apoptotic gene targets as did Lu et al., we did identify REST-bound genes encoding proteins related to preserving brain health. In this group, several of our qChIP-validated genes relating to innate immunity have been implicated in AD (Fig. 3b; Additional file 2: Table S4) and would be predicted to increase with reduced levels of REST [21]. For example, Arachidonate 5-Lipoxygenase *(ALOX5)*, an enzyme that converts arachidonic acid to leukotrienes, is expressed to abnormally high levels in AD in human brains [58]. Interestingly, we also identified Coactosin-like protein *(COTL1)* as a REST target gene. Some evidence supports this protein as an ALOX5 chaperone and increased *COTL1* expression in microglia is associated with AD neuropathology [59]. Expression of the human REST target gene, Diaphanous Related Formin 1 *(DIAPH1)*, involved in actin polymerization, is increased in neurons and glia in AD [60]. A few of our REST target genes also encode proteins involved in brain pathology other than AD. For example, *NLRP4* is involved in formation of the inflammasome complex. Significant enrichment of REST-bound genes in the Interleukin Reactome category (Additional file 2: Table S4) is also indicative of inflammatory processes.

A recent single cell RNA seq analysis of human hippocampus identified REST transcripts in neurons and glia cells (http://dropviz.org/, [61]), consistent with our detection of REST epitopes in the nuclei of both neurons and glia in post mortem hippocampal sections (Additional file 1: Figure S1.3c). Therefore, it’s possible that the REST gene targets we identified in intact hippocampus represented genes in both neuronal and non-neuronal cells types, and that the lower REST peak densities on some target genes may reflect the tissue heterogeneity. The heterogeneity, as well as the age of samples, could also account for the difference in REST target genes identified in this study and the genes expressed specifically in neurons in Lu et al., 2014. Indeed, from published literature, many of the validated REST target genes we identified from our ChIP-seq analysis are expressed in both neurons and glia. Further, a subset of them are expressed normally at fairly low levels but can be induced in response to stimuli. For example, the REST target serum complement genes, C1QA and C1QB, which are expressed to their highest levels specifically in microglia, are still present at minimal levels in healthy brain [62]. However, increased levels of C1Q protein results in AD synapse pathology [63], and this gene induction is compatible with the reduction of REST binding that occurs with AD [21]. We speculate that REST is serving as a chromatin repressor to poise expression of complement genes in hippocampus, and likely other induced innate immunity target genes as well, for future responsiveness to environmental or disease signaling, just as REST serves to poise expression of neuronal genes in progenitor stem cells awaiting the cue to differentiate into neurons [7].

Altogether, including studies indicating an association of human REST single nucleotide polymorphisms with AD [25], accumulating data points to a role for REST levels in preserving a healthy human brain. To more definitively clarify the role for REST in brain pathologies, including aging and AD, REST ChIP-seq analyses, or more sensitive methods, will need be performed on sorted cells from the brain regions, and coordinated with RNA-seq and single cell ATAC analyses [64, 65]. These studies will also open the door to mechanistic studies to address fundamental gaps in our knowledge of repressors generally, such as precisely how REST is recruited to immune and inflammatory genes lacking an RE1 sequence, and, conversely, how REST is prevented from binding to the consensus RE1 sequence in the prototypical gene targets related to neuronal function. Finally, the function of REST in human glia is still ill-defined, but REST repression in these and other non-neuronal cells, such as astrocytes and microglia, likely also contribute importantly to maintaining human brain health.

## Methods

### Mouse strains

*REST^**GT(D047E11)**^* (*Rest^GT^*) mutants were established by blastocyst injection of the D047E11 GT clone (GenBank Acc.: <u>DU821609</u>; [67]). *REST^GTi(D047E11)^* mice carrying the inverted GT vector (*Rest^GTi^*) were obtained by crossing to Flpe deleter mice [68]. This colony is maintained and genotyped to generate brain-specific *REST^/-^* mice using *Nestin Cre* as described in Nechiporuk et al., 2016. Mice were backcrossed to WT C57Bl6 line (Jackson Laboratories Strain 000664) for at least ten generations.

### Human tissue

Tissue was procured through the Oregon Alzheimer’s Disease Center (ADC) and OHSU Department of Pathology and subjects had no significant brain pathologies except age-related changes as established by review of clinical histories and survey of common neurodegenerative disease-associated lesions. Tissues were deidentified and their use was exempted from review by the IRB at OHSU in accordance with relevant guidelines. Informed consent was obtained from all participants and/or their legal representatives. All analyses were performed on postmortem frozen tissue with a time interval from death to preservation of < 24h.

### Immunohistochemistry

Transcardially perfused brains from adult mice were fixed with 4% paraformaldehyde followed by post-fix at 4°C overnight, equilibrated in 30% sucrose and embedded in TFM tissue frozen medium (TBS) for frozen sectioning. Sections were dried, post-fixed with cold acetone and stained using standard immunohistochemical techniques. Antigen retrieval in 10 mM sodium citrate, pH 6.0, 0.1% Tween-20, 95°C, 10 min was used to expose nuclear antigens. Sections were stained with secondary species-specific antibodies conjugated to Alexa-488, Alexa-555, or Alexa-647 (Invitrogen, Waltham, MA), and counterstained with DAPI to reveal nuclei. Immunostaining was analyzed with a confocal fluorescent microscope (Zeiss LSM710 Axiovert). Human tissues were evaluated as described previously [69]. In brief, brains were fixed in neutral-buffered formaldehyde solution for at least 2 weeks and examined grossly and microscopically. For microscopic evaluation, tissue samples were processed into paraffin blocks and 6-micrometer sections were stained with hematoxylin-eosin and Luxol fast blue and immunostained as described above after antigen retrieval with development using ABC kits from Vector Laboratories (Burlingame, CA).

### Antibodies

Chicken anti-MAP2 (Abcam ab5392, RRID:AB_2138153, IF-1:500, Additional file 1: Figure S1.1C), rabbit anti-HDAC2 (Life Technologies, RRID:AB_2547079, WB-1:5000, Additional file 1: Figure S1.2C); Mouse anti-α-tubulin (DSHB clone AA4.3, RRID:AB579793, WB-1:5000, Additional file 1: Figure S1.2D); Rat monoclonal antimouse REST antibody 4A9 (in-house, IF-1:300, Additional file 1: Figure S1.1b,c); Rabbit anti-human REST antibody REST-C [7](in-house, WB-1:1000, Additional file 1: Figures S1.1d, S1.2c, and S1.2d, all used for ChIP); Rabbit anti-human REST (Bethyl A300, RRID:AB_477959, IF-1:500, Additional file 1: Figure S1.2).

### Western Blotting

For human samples, hippocampal tissue was lysed in cold lysis buffer containing 50 mM Tris, pH 7.5, 150 mM NaCl, 1 mM EDTA, 0.1 mM sodium vanadate, 10 mM β-glycerophosphate; 10 mM NaF, protease inhibitors (Roche Basel, Switzerland), 1% Triton X-100 and 10% glycerol and resolved on 3–8% pre-cast gradient Tris-Acetate gels (Invitrogen, Waltham, MA). For mouse samples, nuclear extracts were prepared by the modified Dignam method as previously described [70]. Blotting with primary antibodies was carried out at 4°C overnight followed by incubation with the appropriate secondary antibodies and developed using chemiluminescent West Pico detection kit (Thermo Scientific). Western blot quantification was carried out by subtracting the background signal from the individual bands. REST signal was normalized to loading controls recognized by either HDAC2 or α-tubulin.

### REST ChIP

Approximately 100-200 mg tissue from mouse hippocampus/ human hippocampus was cross-linked in Cross-linking buffer (10 mM HEPES, pH 7.2, 100 mM NaCl, 1 mM EDTA and 1 mM EGTA) containing 1% formaldehyde (Thermo; methanol free) for 10 minutes at room temperature. Excess formaldehyde was quenched with 125 mM glycine for 5 minutes at room temperature. Tissue was resuspended in homogenization buffer (250 mM Sucrose, 25 mM KCl, 5 mM MgCl_2_, 20 mM Tricine-KOH; pH 7.8, 1 mM DTT, 0.15 mM spermine, 0.5 mM spermidine, and protease inhibitors) after a wash with ice-cold PBS. Tissue was homogenized using Dounce homogenizer with loose (A) and tight (B) pestle (5 strokes each), followed by additional 5 strokes of tight pestle with 0.3% NP-40, and then the homogenate was passed through a 40 *μ*m strainer. Nuclei were isolated at 4000g/ 4°C for 5 minutes and resuspended in ~0.5 ml sonication buffer (10 mM Tris-HCl pH 8.0, 1 mM EDTA, 1 mM EGTA). Fragmented chromatin was prepared from ~10 million nuclei in Covaris S220 (200 cycle per burst, 5% duty cycle at power level 4 for 12 minutes at 4°C) in presence of 0.3% SDS. For ChIP, Protein G Dynabeads (Invitrogen; 30 *μ*l/reaction) were equilibrated in TBSTBp (TBS with 1% BSA, 1% Triton-X-100, and protease inhibitors). 10 *μ*l Protein G beads were used for preclearing the chromatin in ChIP buffer (10 mM Tris-HCl pH 8.0, 1% Triton-X-100, 150 mM NaCl and 1 mM EDTA, and protease inhibitors) for 4 hours at 4°C. 20 *μ*l Protein G beads were used to make the bead-Ab complex with 20 *μ*g anti-REST-C antibodies [7] for 2 hours at room temperature in 0.5 ml TBSTBp. After washing bead-Ab complex three times with TBSTBp, ChIP was carried out overnight at 4°C with precleared chromatin. Beads were washed twice with low salt, high salt and LiCl buffers at 4°C, followed by two washes with TE at room temperature. DNA was eluted and crosslinks were reversed overnight at 65°C. Contaminants were removed from DNA by RNaseA and Proteinase K treatment followed by purification with PCR purification kit (QIAGEN).

### REST ChIP-seq

Ten nanograms of fragmented DNA was used as input for a modified TruSeq Nano DNA library preparation protocol (Illumina). Briefly, input DNA was treated with 3’ to 5’ exonuclease activity and 5’ to 3’ polymerase activity to blunt the ends. There was no size selection of the fragments. A single “A” nucleotide was added to the 3’ ends to enhance ligation to the adapters. RNA adapters (Illumina) were ligated to the fragmented DNA followed by cleanup using sample purification beads (SPB) provided with the kit. The ligation product was enriched using 14 cycles of polymerase chain reaction. The amplification product was cleaned using SPB. Libraries were profiled using the TapeStation D1000 DNA tape. Library concentrations were determined using the Library Quantification Kit for Illumina sequencing platforms (Kapa Biosystems) on a StepOnePlus Real Time PCR Workstation (ThermoFisher). Libraries were mixed for multiplexing and the concentration of the mix was determined by real time PCR. Short read sequencing was performed on a HiSeq 2500 (Illumina) using a single 100-cycle protocol by the OHSU Massively Parallel Sequencing Shared Resource. Base call files were converted to fastq files using Bcl2Fastq2 (Illumina).

### ChIP-seq analysis

Sequenced reads were processed and analyzed by the ENCODE ChIP-seq analysis pipeline (https://github.com/ENCODE-DCC/chip-seq-pipeline2) using the GRCh38 (hg38) version of the human genome and the mm10 version of the mouse genome. Three human replicates (age 63, 63, and 67) were processed. For mice, two replicates of pooled hippocampi from 4 animals were analyzed. Peaks for each replicate were called and then the final peak list was based on the optimal set calculated using the SPP peak caller and the irreproducible discovery rate (IDR), which incorporates all the replicates into the analysis.

### Read quality analysis, gene annotation, and Gene Ontology enrichment

REST peaks for each experiment were converted into bed files and analyzed using the R programs Homer [71], ChIPseeker [72] and ClusterProfiler [73]. REST target genes associated with all peaks used for Gene Ontology analysis were annotated with the seq2gene function in ChIPseeker. REST target genes were defined as genes with REST peaks located within the gene body and including up to 5kb upstream of the Refseq-annotated transcriptional start site (TSS). ‘Core’ REST peaks were defined as peaks that overlapped in at least 40 datasets of the 48 non-neuronal REST ChIP-seq replicates publicly available from ENCODE as determined by Homer (Additional file 2: Table S5). Gene Ontology lists were reduced in complexity by the PANTHER ‘slim’ database [74], and overrepresentation and enrichment of GO terms was calculated using Fisher’s exact test with Bonferroni correction. Motif analysis was performed using DREME [75] and DIVERSITY [34] on all REST peaks from mouse or human hippocampus.

### HOT region filtering

As known HOT regions predominately centered around highly expressed TSS [76], we used aggregate plots for ChIP-seq read depths for H3K4me3 and H3K9ac modifications to simulate HOT regions in hippocampal tissue and the cell lines analyzed. We then calculated an active chromatin score for each putative HOT region and compared them to REST enrichment levels. We first tested our filter method on two independent ENCODE datasets from human Embryonic Stem Cells (ESCs) and the human A459 lung carcinoma cell line (Additional file 2: Table S5), and were able to establish a threshold that eliminated the majority of the TSS H3K4me3 and H3K9ac peaks, 63% and 79% respectively, while maintaining the majority, 87% and 99%, of REST peaks containing RE1 sites (Additional file 3: Figure S1c). We then extended the filter to ENCODE ChIP-seq analysis on human H1 neurons and our human hippocampal dataset, normalizing the thresholds to median REST and H3K4me3 and H3K9ac ChIP-seq reads for each dataset. The filter significantly reduced the proportion of HOT regions with minimal loss of REST peaks in all of these cell types and human hippocampus, and, importantly, in hippocampal peaks associated with REST RE1 sequences (Additional file 3: Figure S1d, e). As predicted, the filtered peaks centered on promoter regions and the closest annotated genes to the filtered peaks were enriched significantly for housekeeping functions atypical of REST target genes and typical of HOT regions, such as ribosome biogenesis, splicing, and DNA metabolism. In total, our filter removed 11,495 of 13,620 REST peaks.

## Supplementary information

*Additional file 2: Table S1. REST ChIP-seq peaks and associated genes in mouse hippocampus*.

*Additional file 2: Table S2. GO and Reactome analysis of mouse REST target genes in hippocampus, mESC and mC2C12 datasets*.

*Additional file 2: Table S3. REST ChIP-seq peaks and associated peaks in human hippocampus*.

*Additional file 2: Table S4. GO and Reactome analysis of human REST target genes in hippocampus and genes overlapping Core*.

*Additional file 2: Table S5. All additional sources for datasets used for analysis*.

## Acknowledgments

The authors thank all members of the Mandel lab for discussion and support, staff at the OHSU MPSSR for their expertise, and Dr. Andrew Adey (OHSU) for critical comments on the bioinformatics analyses.

## Authors’ contributions

G.M. J.M. M.S, S.G, and K.M. designed the research. S.G, M.S, K.M, and R.W. generated the data. J.M performed the bioinformatics analysis. G.M. wrote the manuscript, with contributions from all of the co-authors. All authors read and approved the final manuscript.

## Data access

ChIP-seq datasets generated in the current study have been deposited in the NCBI Gene Expression Omnibus (GEO) database (https://www.ncbi.nlm.nih.gov/geo/) and are accessible through accession number GSE144226. Previously published datasets used for comparative analysis are listed and cited in Additional file 2: Table S5.

## Ethics approval

All mouse procedures were approved by the Institutional Animal Care and Use Committee at OHSU (IACUC No. IP00000284) and carried out in accordance with approved guidelines.

---

**Additional file 1: Figure S1.1.**
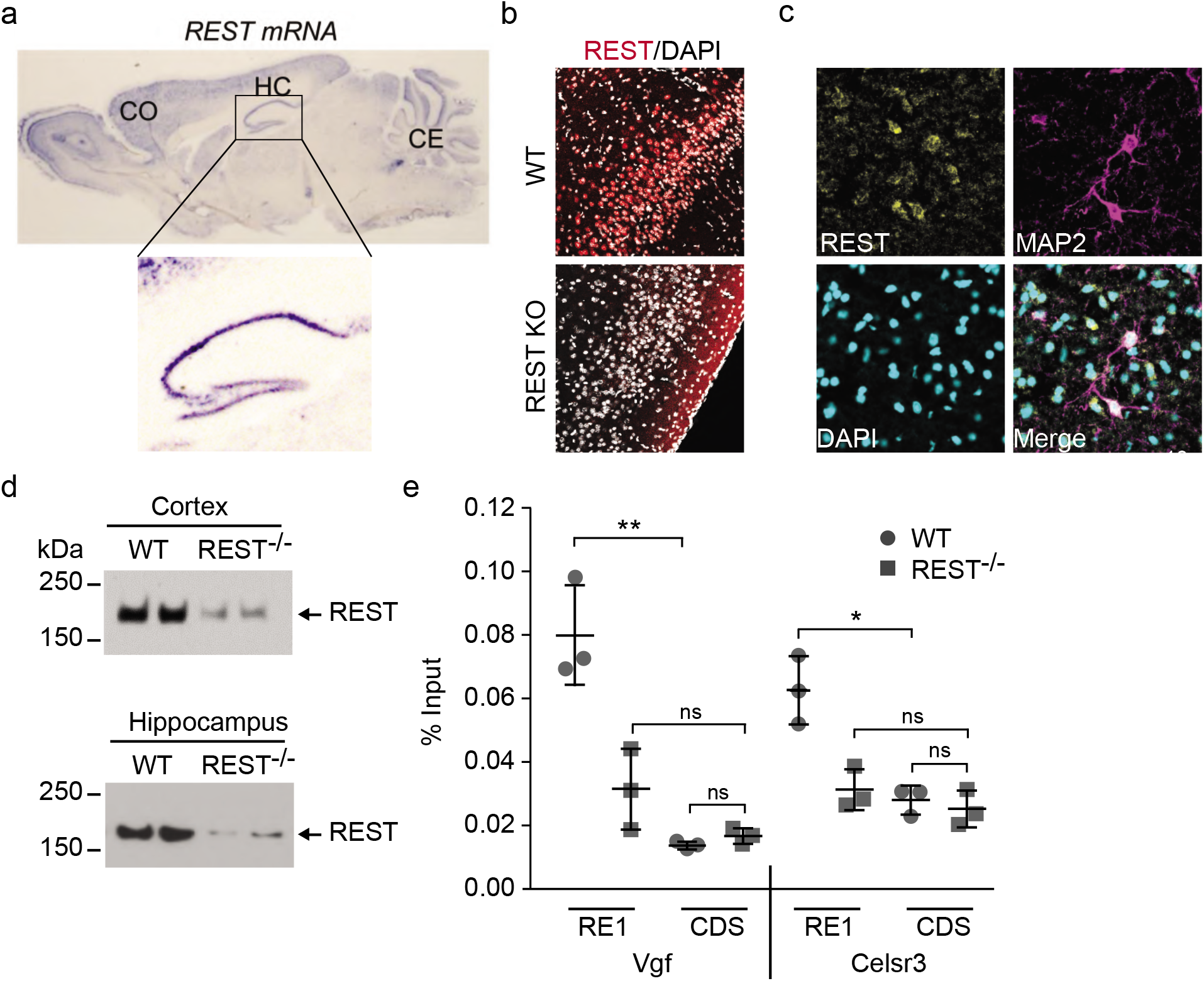
REST expression in hippocampus and validation of antibody specificity in REST knockout mouse brain. a) In situ hybridization for REST transcripts throughout the postnatal brain of 5-week old male mouse. Inset shows hippocampal in situ hybridization signal in the molecular layer. b) REST immunoreactivity in wild type juvenile (P30) mouse piriform cortex is greatly reduced in REST neural-progenitor-specific (Nestin) knockout. DAPI, nuclear marker. c) REST immunoreactivity in juvenile cortex (P30) coincides with MAP2-positive neurons as well as MAP2-negative cells. DAPI identifies nuclei. d) Western blot showing reduction of REST protein (~200 kDa) in two brain regions isolated from wild type (WT) and neural progenitor (Nestin)-specific REST knockout mice. n=1 mouse/lane from 5w old male mice. e) Quantitative ChIP results for REST binding in hippocampus of 5w old male mice in the REST neural progenitor-specific knockout samples compared to wild type samples. Coding sequences (CDS) in indicated genes were used as controls.

**Additional file 1: Figure S1.2.**
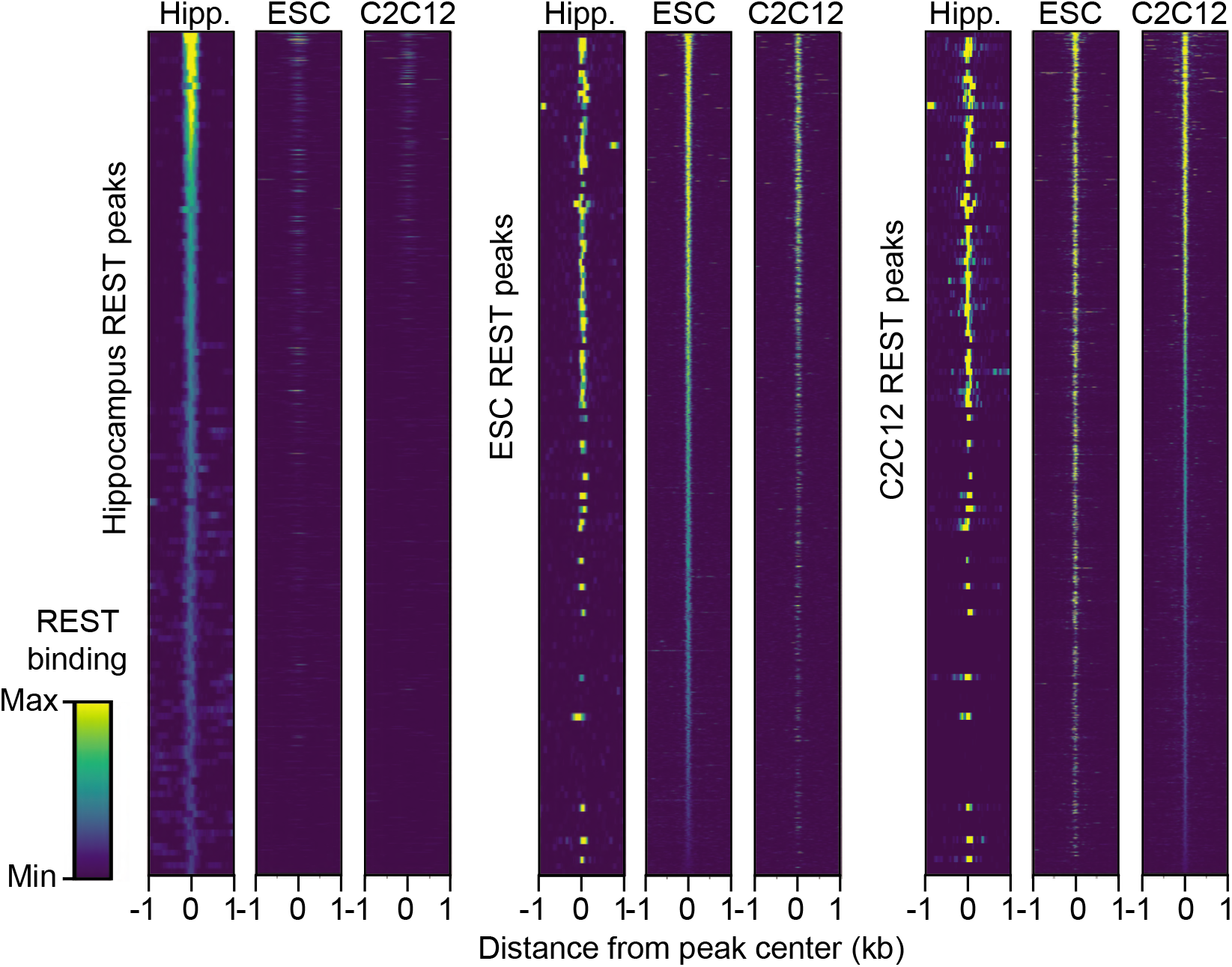
REST densities in REST peaks in mouse brain. Heat maps of REST binding density at ChIP-seq peaks. Each heat map is scaled to the maximum level of REST bound in each tissue/cell type, and regions are sorted from highest to lowest REST binding levels in hippocampus (left), embryonic stem cells (center) and myoblast C2C12 cells (right).

**Additional file 1: Figure S1.3.**
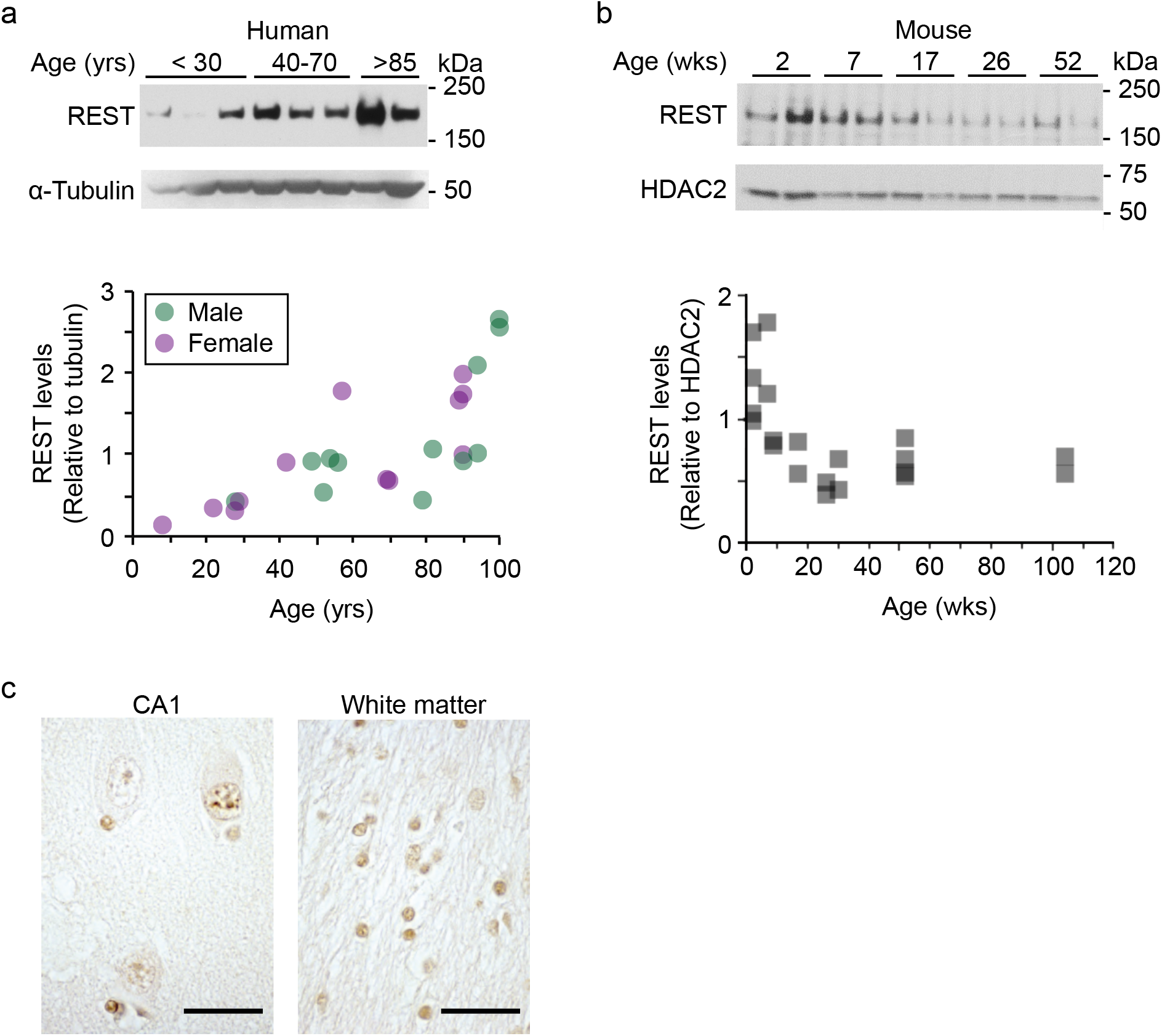
REST protein levels in human and mouse hippocampal extracts and in human hippocampus sections. a) (top) Representative Western blot of human hippocampal REST protein levels in extracts prepared from female samples of indicated ages. Proteins were fractionated by polyacrylamide gel electrophoresis and membranes were probed for REST and a-tubulin protein (bottom). Quantification of aggregate Western blot data including samples in top Western blot (12 females, purple circles; 12 males, green circles). a-tubulin is abundant and levels are unchanged with age. b) (top) As in a) but Western blot used male mouse hippocampal extracts and membranes were probed for REST and HDAC2. (Bottom) Quantification of aggregate Western blot data (20 males including data from top Western blot). Note variations in HDAC2 intensity that we assume are due to variations in sample loading as HDAC2 levels remain consistent with postnatal age [66], REST migrates at ~200kDa. c) Immuno-labeling of human hippocampal pyramidal and granular sectors (left) and oligodendroglial temporal white matter (right) shows nuclear staining of REST with an affinity-purified REST antibody made in-house (anti-C terminal [15]). Higher magnification staining shows REST in both neurons and glia that is almost exclusively nuclear. Scale bar = 30 μm

**Additional file 1: Figure S2.1.**
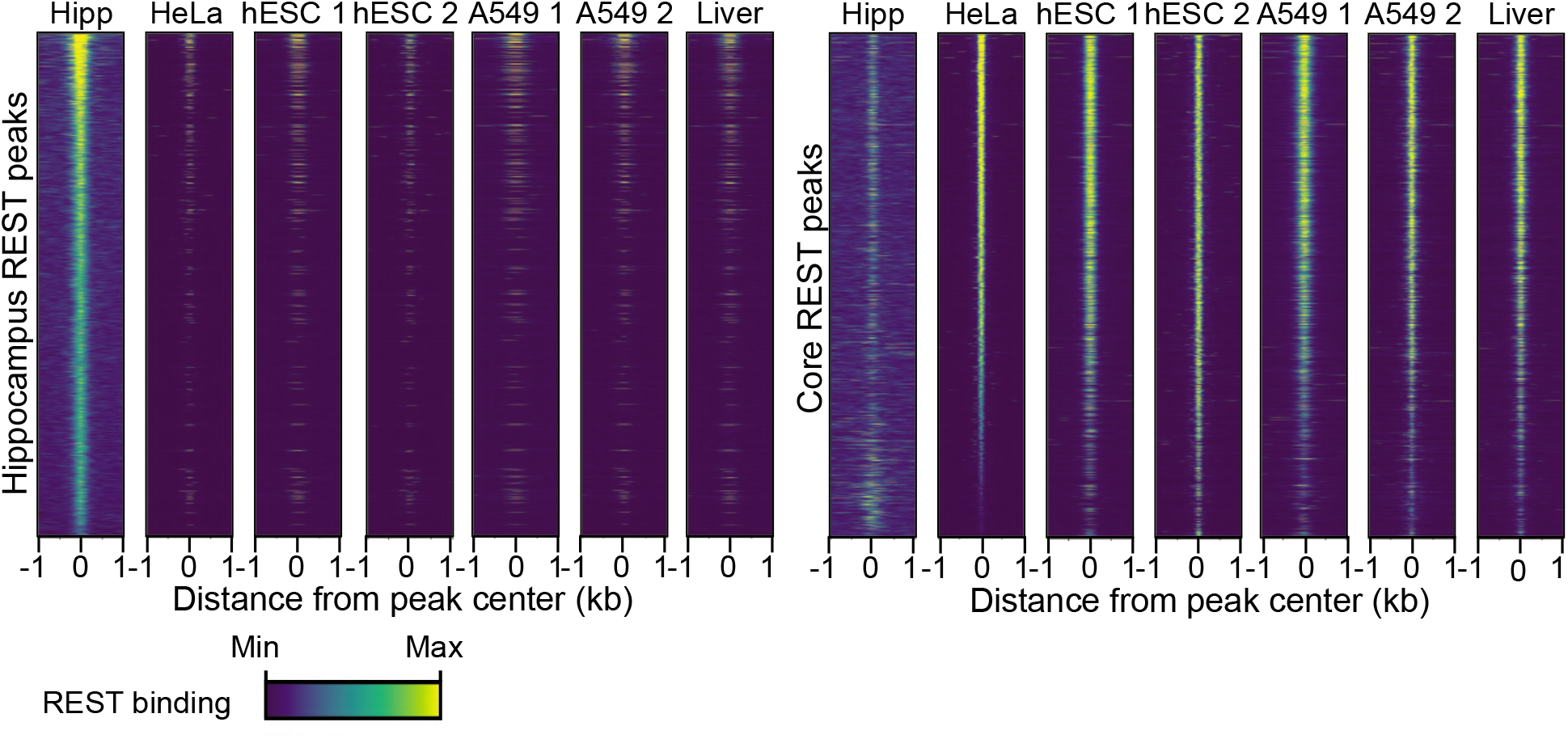
Human hippocampal REST ChIP-seq data is distinct from other REST ChIP-seq datasets. Heap maps representing the density of cell type-specific REST binding at peaks from our human hippocampal ChIP-seq dataset (left) or from the Core ENCODE ChIP-seq dataset (right). Each heatmap is scaled to the maximum level of REST bound in each tissue/cell type.

**Additional file 1: Figure S3.1.**
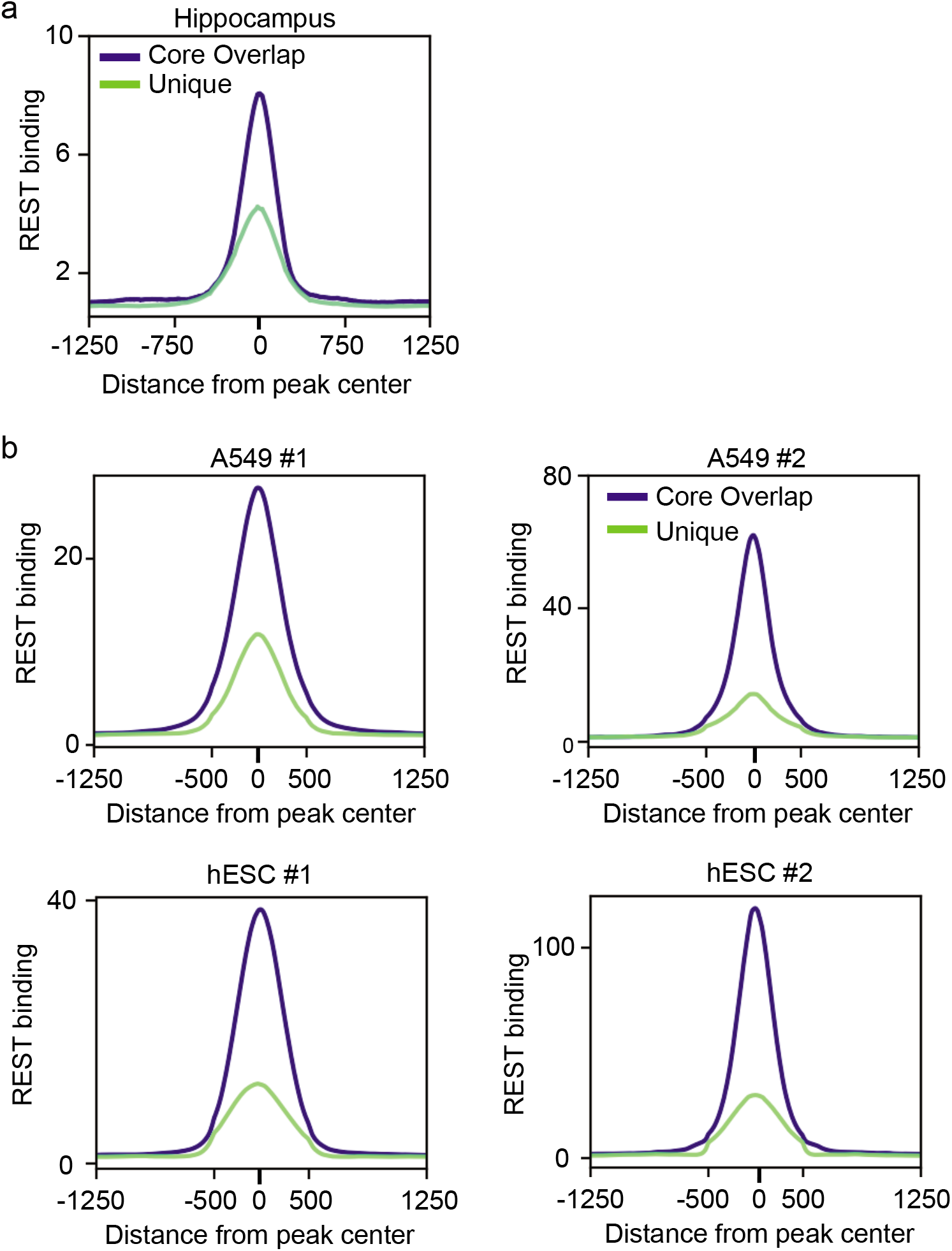
Amount of REST bound at peaks unique to tissue and cell type. a) Amount of REST bound at hippocampal unique REST peaks (green) is lower than that observed at loci shared with Core REST peaks (blue). Core as defined in Figure 2. b) Mean intensity plots for REST in REST peaks for A549 and hESC cells compared to other cell types in ENCODE, parsed by peaks unique to ESC or A549 (green) or peaks that overlap Core peaks (blue).

**Additional file 3: Figure S1.**
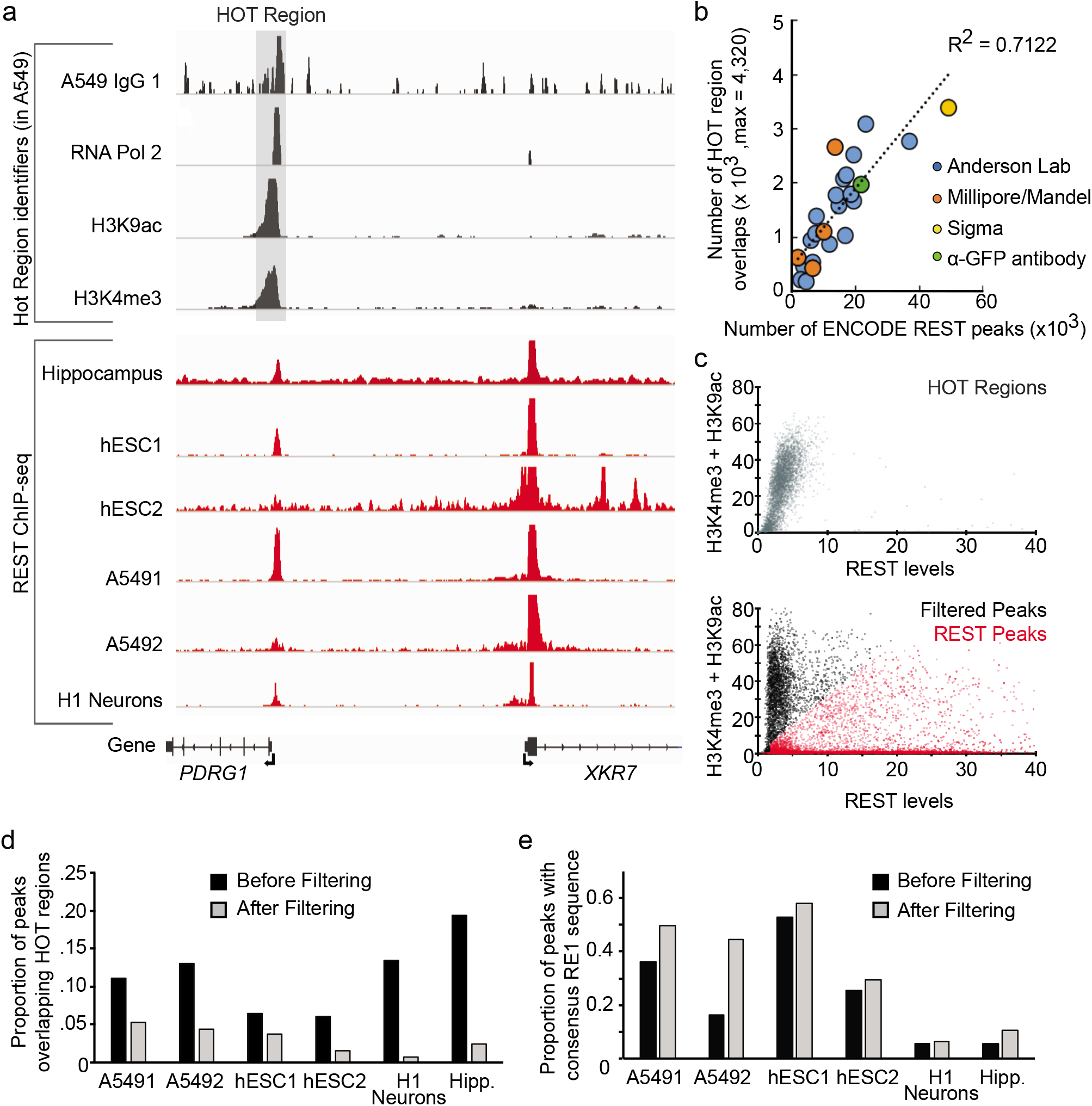
Bioinformatics filtering of REST ChIP-seq diminishes HOT regions while preserving most REST peaks. a) (Top traces) Representative ChIP-seq analysis for TSSs as defined by ChIP-seq analyses (ENCODE) for RNA polymerase II and active transcription markers (H3K9Ac; H3K4me3) in the A549 cell line (proxy for HOT regions). Note HOT region even in the IgG1 ChIP control trace (ENCFF593HFQ). (Bottom traces) Our REST ChIP-seq analysis in indicated cell lines and human H1 neurons in the same genomic regions as in a). Note presence of REST peaks that overlap with HOT regions. b) Scatter plot of REST ChIP-seq peaks (ENCODE) using different antibody sources and their correlation with annotated HOT regions [35]. c) Scatter plots of REST ChIP-seq data from the A549 cell line at annotated HOT regions (Top) or all REST peaks (Bottom), color-coded for those that pass the filter (red) and those that fail (black). d and e) Histograms of the proportion REST ChIP-seq peaks that remain after filtering in indicated cells lines and H1 neurons. (d) Proportion of total REST peaks that overlap HOT regions and e) proportion of REST peaks with underlying consensus RE1 sequences.

## References

1. Chong JA, Tapia-Ramírez J, Kim S, Toledo-Aral JJ, Zheng Y, Boutros, MC, Altshuller YM, Frohman MA, Kraner SD, and Mandel G. REST: a mammalian silencer protein that restricts sodium channel gene expression to neurons. Cell. 1995;80:949–957.

2. Schoenherr CJ, and Anderson DJ. The neuron-restrictive silencer factor (NRSF): a coordinate repressor of multiple neuron-specific genes. Science. 1995;267:1360–1363.

3. Kraner SD, Chong JA, Tsay H-J, and Mandel G. Silencing the type II sodium channel gene: A model for neural-specific gene regulation. Neuron. 1992;9:37–44.

4. Schoenherr CJ, Paquette AJ, Anderson DJ. Identification of potential target genes for the neuron-restrictive silencer factor. Proc Natl Acad Sci U S A. 1996;93(18):9881–9886.

5. Otto SJ, McCorkle SR, Hover J, Conaco C, Han J-J, Impey S, Yochum GS, Dunn JJ, Goodman RH, and Mandel G. A new binding motif for the transcriptional repressor REST uncovers large gene networks devoted to neuronal functions. J. Neurosci. 2007;27:6729–6739.

6. Johnson DS, Mortazavi A, Myers RM, and Wold B. Genome-wide mapping of in-vivo protein-DNA interactions. Science. 2007;316:1497–1502.

7. Ballas N, Grunseich C, Lu DD, Speh JC, and Mandel G. REST and its corepressors mediate plasticity of neuronal gene chromatin throughout neurogenesis. Cell. 2005;121:645–657.

8. Conaco C, Otto S, Han JJ, Mandel G. Reciprocal actions of REST and a microRNA promote neuronal identity. Proc Natl Acad Sci U S A. 2006;103(7):2422–2427.

9. Ballas N, Mandel G. The many faces of REST oversee epigenetic programming of neuronal genes. Curr Opin Neurobiol. 2005;15(5):500–506.

10. Palm K, Belluardo N, Metsis M, Timmusk T. Neuronal expression of zinc finger transcription factor REST/NRSF/XBR gene. J Neurosci. 1998;18(4):1280–1296.

11. Gao Z, Ure K, Ding P, et al. The master negative regulator REST/NRSF controls adult neurogenesis by restraining the neurogenic program in quiescent stem cells. J Neurosci. 2011;31 (26):9772–9786.

12. Calderone A, Jover T, Noh KM, et al. Ischemic insults derepress the gene silencer REST in neurons destined to die. J Neurosci. 2003;23(6):2112–2121.

13. Kuwabara T, Hsieh J, Nakashima K, Taira K, Gage FH. A small modulatory dsRNA specifies the fate of adult neural stem cells. Cell. 2004;116(6):779–793.

14. Sun YM, Greenway DJ, Johnson R, et al. Distinct profiles of REST interactions with its target genes at different stages of neuronal development [published correction appears in Mol Biol Cell. 2006 Mar;17(3):1494]. Mol Biol Cell. 2005;16(12):5630–5638.

15. Nechiporuk T, McGann J, Mullendorff K, et al. The REST remodeling complex protects genomic integrity during embryonic neurogenesis. Elife. 2016;5:e09584.

16. Noh KM, Hwang JY, Follenzi A, et al. Repressor element-1 silencing transcription factor (REST)-dependent epigenetic remodeling is critical to ischemia-induced neuronal death. Proc Natl Acad Sci U S A. 2012;109(16):E962–E971.

17. Kaneko N, Hwang JY, Gertner M, Pontarelli F, Zukin RS. Casein kinase 1 suppresses activation of REST in insulted hippocampal neurons and halts ischemia-induced neuronal death. J Neurosci. 2014;34(17):6030–6039.

18. Hu XL, Cheng X, Cai L, et al. Conditional deletion of NRSF in forebrain neurons accelerates epileptogenesis in the kindling model. Cereb Cortex. 2011;21(9):2158–2165.

19. McClelland S, Brennan GP, Dubé C, et al. The transcription factor NRSF contributes to epileptogenesis by selective repression of a subset of target genes. Elife. 2014;3:e01267.

20. Singh-Taylor A, Molet J, Jiang S, et al. NRSF-dependent epigenetic mechanisms contribute to programming of stress-sensitive neurons by neonatal experience, promoting resilience. Mol Psychiatry. 2018;23(3):648–657.

21. Lu T, Aron L, Zullo J, et al. REST and stress resistance in ageing and Alzheimer’s disease. Nature. 2014;507(7493):448–454.

22. Zullo JM, Drake D, Aron L, et al. Regulation of lifespan by neural excitation and REST. Nature. 2019;574(7778):359–364.

23. Mortazavi A, Leeper Thompson EC, Garcia ST, Myers RM, Wold B. Comparative genomics modeling of the NRSF/REST repressor network: from single conserved sites to genome-wide repertoire. Genome Res. 2006;16(10):1208–1221

24. Aigner S, and Yeo GW. Terminal Differentiation: REST. In: Squire LR, editor. Encyclopedia of Neuroscience. Oxford: Academic Press; 2009. p. 921–927.

25. Mozzi A, Guerini FR, Forni D, et al. REST, a master regulator of neurogenesis, evolved under strong positive selection in humans and in non human primates. Sci Rep. 2017;7(1):9530.

26. Rockowitz S, Lien WH, Pedrosa E, et al. Comparison of REST cistromes across human cell types reveals common and context-specific functions. PLoS Comput Biol. 2014;10(6):e1003671.

27. Rockowitz S, Zheng D. Significant expansion of the REST/NRSF cistrome in human versus mouse embryonic stem cells: potential implications for neural development. Nucleic Acids Res. 2015;43(12):5730–5743.

28. Wang D, Liu S, Warrell J, et al. Comprehensive functional genomic resource and integrative model for the human brain. Science. 2018;362(6420):eaat8464.

29. Johnson R, Samuel J, Ng CK, Jauch R, Stanton LW, Wood IC. Evolution of the vertebrate gene regulatory network controlled by the transcriptional repressor REST. Mol Biol Evol. 2009;26(7):1491–1507.

30. McGann JC, Oyer JA, Garg S, et al. Polycomb-and REST-associated histone deacetylases are independent pathways toward a mature neuronal phenotype. Elife. 2014;3:e04235.

31. Davis CA, Hitz BC, Sloan CA, et al. The Encyclopedia of DNA elements (ENCODE): data portal update. Nucleic Acids Res. 2018;46(D1):D794–D801.

32. Chen GL, Miller GM. Alternative *REST* Splicing Underappreciated. eNeuro. 2018;5(5):ENEURO.0034–18.2018.

33. He P & Williams BA, Trout D, et al. The changing mouse embryo transcriptome at whole tissue and single-cell resolution. bioRxiv 2020;150599. doi:10.1101/2020.06.14.150599

34. Mitra S, Biswas A, Narlikar L. DIVERSITY in binding, regulation, and evolution revealed from high-throughput ChIP. PLoS Comput Biol. 2018;14(4):e1006090.

35. Wreczycka K, Franke V, Uyar B, et al. HOT or not: examining the basis of high-occupancy target regions. Nucleic Acids Res. 2019;47(11):5735–5745.

36. Park D, Lee Y, Bhupindersingh G, Iyer VR. Widespread misinterpretable ChIP-seq bias in yeast. PLoS One. 2013;8(12):e83506.

37. Teytelman L, Thurtle DM, Rine J, van Oudenaarden A. Highly expressed loci are vulnerable to misleading ChIP localization of multiple unrelated proteins. Proc Natl Acad Sci U S A. 2013;110(46): 18602–18607.

38. Jain D, Baldi S, Zabel A, Straub T, Becker PB. Active promoters give rise to false positive ‘Phantom Peaks’ in ChIP-seq experiments. Nucleic Acids Res. 2015;43(14):6959–6968.

39. Krebs W, Schmidt SV, Goren A, et al. Optimization of transcription factor binding map accuracy utilizing knockout-mouse models. Nucleic Acids Res. 2014;42(21):13051–13060.

40. Bruce AW, Donaldson IJ, Wood IC, et al. Genome-wide analysis of repressor element 1 silencing transcription factor/neuron-restrictive silencing factor (REST/NRSF) target genes. Proc Natl Acad Sci U S A. 2004;101(28):10458–10463.

41. Anazi S, Maddirevula S, Salpietro V, et al. Expanding the genetic heterogeneity of intellectual disability [published correction appears in Hum Genet. 2017 Dec 29;:]. Hum Genet. 2017;136(11-12):1419–1429.

42. Fabregat A, Jupe S, Matthews L, et al. The Reactome Pathway Knowledgebase. Nucleic Acids Res. 2018;46(D1):D649–D655.

43. Gupta S, Stamatoyannopoulos JA, Bailey TL, Noble WS. Quantifying similarity between motifs. Genome Biol. 2007;8(2):R24.

44. Gupta P, Gurudutta GU, Saluja D, Tripathi RP. PU.1 and partners: regulation of haematopoietic stem cell fate in normal and malignant haematopoiesis. J Cell Mol Med. 2009;13(11-12):4349–4363.

45. Chang DH, Angelin-Duclos C, Calame K. BLIMP-1: trigger for differentiation of myeloid lineage. Nat Immunol. 2000;1(2):169–176.

46. Tamura T, Yanai H, Savitsky D, Taniguchi T. The IRF family transcription factors in immunity and oncogenesis. Annu Rev Immunol. 2008;26:535–584.

47. Wang J, Dayyani F, Brynzka C, Sweetser D. TLEs regulate myeloid proliferation and differentiation in association with modulation of NF-κB and Wnt signaling. Blood. 2010;116(21):4176.

48. Terrados G, Finkernagel F, Stielow B, et al. Genome-wide localization and expression profiling establish Sp2 as a sequence-specific transcription factor regulating vitally important genes. Nucleic Acids Res. 2012;40(16):7844–7857.

49. Growney JD, Shigematsu H, Li Z, et al. Loss of Runx1 perturbs adult hematopoiesis and is associated with a myeloproliferative phenotype. Blood. 2005;106(2):494–504.

50. van der Meer LT, Jansen JH, van der Reijden BA. Gfi1 and Gfi1b: key regulators of hematopoiesis. Leukemia. 2010;24(11):W34–1843.

51. Andrés ME, Burger C, Peral-Rubio MJ, et al. CoREST: a functional corepressor required for regulation of neural-specific gene expression. Proc Natl Acad Sci U S A. 1999;96(17):9873–9878.

52. Gu X, Hu Z, Ebrahem Q, et al. Runx1 regulation of Pu.1 corepressor/coactivator exchange identifies specific molecular targets for leukemia differentiation therapy. J Biol Chem. 2014;289(21):14881–14895.

53. Yamamoto R, Kawahara M, Ito S, et al. Selective dissociation between LSD1 and GFI1B by a LSD1 inhibitor NCD38 induces the activation of *ERG* super-enhancer in erythroleukemia cells. Oncotarget. 2018;9(30):21007–21021.

54. Roth M, Bonev B, Lindsay J, et al. FoxG1 and TLE2 act cooperatively to regulate ventral telencephalon formation. Development. 2010;137(9):1553–1562.

55. Mukherjee S, Brulet R, Zhang L, Hsieh J. REST regulation of gene networks in adult neural stem cells. Nat Commun. 2016;7:13360.

56. Meers MP, Bryson TD, Henikoff JG, Henikoff S. Improved CUT&RUN chromatin profiling tools. Elife. 2019;8:e46314.

57. Ramani V, Qiu R, Shendure J. High Sensitivity Profiling of Chromatin Structure by MNase-SSP. Cell Rep. 2019;26(9):2465–2476.e4.

58. Firuzi O, Zhuo J, Chinnici CM, Wisniewski T, Praticò D. 5-Lipoxygenase gene disruption reduces amyloid-beta pathology in a mouse model of Alzheimer’s disease. FASEB J. 2008;22(4):1169–1178.

59. Rangaraju, S., Dammer, E.B., Raza, S. et al. Quantitative proteomics of acutely-isolated mouse microglia identifies novel immune Alzheimer’s disease-related proteins. Mol Neurodegeneration. 2018;13(34)

60. Derk J, Bermudez Hernandez K, Rodriguez M, et al. Diaphanous 1 (DIAPH1) is Highly Expressed in the Aged Human Medial Temporal Cortex and Upregulated in Myeloid Cells During Alzheimer’s Disease. J Alzheimers Dis. 2018;64(3):995–1007.

61. Saunders A, Macosko EZ, Wysoker A, et al. Molecular Diversity and Specializations among the Cells of the Adult Mouse Brain. Cell. 2018;174(4):1015–1030.e16.

62. Morgan BP. Complement in the pathogenesis of Alzheimer’s disease. Semin Immunopathol. 2018;40(1):113–124.

63. Hong S, Beja-Glasser VF, Nfonoyim BM, et al. Complement and microglia mediate early synapse loss in Alzheimer mouse models. Science. 2016;352(6286):712–716.

64. Xu X, Stoyanova EI, Lemiesz AE, Xing J, Mash DC, Heintz N. Species and cell-type properties of classically defined human and rodent neurons and glia. Elife. 2018;7:e37551.

65. Nativio R, Donahue G, Berson A, et al. Dysregulation of the epigenetic landscape of normal aging in Alzheimer’s disease [published correction appears in Nat Neurosci. 2018 Mar 19;:]. Nat Neurosci. 2018;21(4):497–505.

66. MacDonald JL, Roskams AJ. Histone deacetylases 1 and 2 are expressed at distinct stages of neuro-glial development. Dev Dyn. 2008;237(8):2256–2267.

67. Hansen J, Floss T, Van Sloun P, et al. A large-scale, gene-driven mutagenesis approach for the functional analysis of the mouse genome. Proc Natl Acad Sci U S A. 2003;100(17):9918–9922.

68. Rodríguez CI, Buchholz F, Galloway J, et al. High-efficiency deleter mice show that FLPe is an alternative to Cre-loxP. Nat Genet. 2000;25(2):139–140.

69. Nguyen MT, Mattek N, Woltjer R, et al. Pathologies Underlying Longitudinal Cognitive Decline in the Oldest Old. Alzheimer Dis Assoc Disord. 2018;32(4):265–269.

70. Grimes JA, Nielsen SJ, Battaglioli E, et al. The co-repressor mSin3A is a functional component of the REST-CoREST repressor complex. J Biol Chem. 2000;275(13):9461–9467.

71. Heinz S, Benner C, Spann N, et al. Simple combinations of lineage-determining transcription factors prime cis-regulatory elements required for macrophage and B cell identities. Mol Cell. 2010;38(4):576–589.

72. Yu G, Wang LG, He QY. ChIPseeker: an R/Bioconductor package for ChIP peak annotation, comparison and visualization. Bioinformatics. 2015;31(14):2382–2383.

73. Yu G, Wang LG, Han Y, He QY. clusterProfiler: an R package for comparing biological themes among gene clusters. OMICS. 2012;16(5):284–287.

74. Mi H, Muruganujan A, Ebert D, Huang X, Thomas PD. PANTHER version 14: more genomes, a new PANTHER GO-slim and improvements in enrichment analysis tools. Nucleic Acids Res. 2019;47(D1):D419–D426.

75. Bailey TL. DREME: motif discovery in transcription factor ChIP-seq data. Bioinformatics. 2011;27(12):1653–1659.

76. Chen RA, Stempor P, Down TA, Zeiser E, Feuer SK, Ahringer J. Extreme HOT regions are CpG-dense promoters in C. elegans and humans. Genome Res. 2014;24(7):1138–1146.

